# DRDs and Brain Derived Neurotrophic Factor share a common therapeutic ground: A novel bioinformatic approach sheds new light towards pharmacological treatment of Cognitive and Behavioral Disorders

**DOI:** 10.1101/2022.09.16.508267

**Authors:** Louis Papageorgiou, Efstathia Kalospyrou, Eleni Papakonstantinou, Io Diakou, Katerina Pierouli, Konstantina Dragoumani, Flora Bacopoulou, George P Chrousos, Themis P Exarchos, Panagiotis Vlamos, Elias Eliopoulos, Dimitrios Vlachakis

## Abstract

Cognitive and behavioral disorders are subgroups of mental health disorders. Both cognitive and behavioral disorders can occur in people of different ages, genders, and social backgrounds and they can cause serious physical, mental or social problems. The risk factors for these diseases are numerous, with a range from genetic and epigenetic factors to physical factors. In most cases, the appearance of such a disorder in an individual is a combination of his genetic profile and environmental stimuli. To date, researchers have not been able to identify the specific causes of these disorders and as such, there is urgent need for innovative study approaches. The aim of the present study was to identify the genetic factors which seem to be more directly responsible for the occurrence of a cognitive and/or behavioral disorder. More specifically, through bioinformatics tools and software as well as analytical methods such as systemic data and text mining, semantic analysis, and scoring functions, we extracted the most relevant single nucleotide polymorphisms (SNPs) and genes connected to these disorders. All the extracted SNPs were filtered, annotated, classified, and evaluated in order to create the “genomic grammar” of these diseases. The identified SNPs guided the search for top suspected genetic factors, dopamine receptors D and Neurotrophic Factor BDNF, for which regulatory networks were built. The identification of the “genomic grammar” and underlying factors connected to cognitive and behavioral disorders can aid the successful disease profiling, the establishment of novel pharmacological targets and provide the basis for personalized medicine, which takes into account the patient’s genetic background as well as epigenetic factors.

## Introduction

Cognitive and Behavioral Disorders are disorders that affect cognition, memory, learning and reading abilities, as well as behavior and movement. These disorders are part of the greater group of mental disorders (1). They can affect people regardless of factors such as age, race or socio-economic status. In cognitive disorders, the grouping includes delirium, mild neurocognitive disorder (also termed mild cognitive disorder), amnestic disorder, and dementia (1). In behavioral disorders, the grouping includes anxiety, depressive, eating, personality, sleep, psychotic, mood and drug and alcohol induced disorders (2). Cognitive disorders affect mostly elderly people, as their developmental phase is connected to rising age. There are currently more than 55 million people living with dementia worldwide and there are almost 10 million new cases each year. Alzheimer’s disease is the most common form of dementia and can contribute to 60-70% of cases (3). In addition, dementia does not exclusively affect the elderly; young dementia, defined as a condition with symptoms onset before the age of 65, accounts for up to 9% of cases (3). On the contrary, behavioral disorders affect individuals, regardless of age (4). According to the World Health Organization (WHO), mental health and behavioral disorders are one of the leading causes of disability worldwide. One in four families is likely to have, in addition to their children, if any, at least one member with a behavioral or mental disorder (2). Statistics from WHO have identified some age groups that are most affected by behavioral disorders: mood disorders (29-34 years), anxiety disorders (24-50 years), substance abuse disorders (18-29 years) (5). However, there is also a high degree of comorbidity, especially in young children or younger adults with mental disorders; of the children diagnosed with a disorder, about 25% appear to have a second one. The percentage of children with comorbidity increases approximately 1.6 times for each additional year from the age of 2 years (18.2%) to 5 (49.7%) (6). Nevertheless, both statistical data from cognitive and behavioral disorders have shown that women are more influenced from these disorders than men (7).

To date, it is known that both an individual’s genetic background and the interacting environment constitute the basic stimuli for the occurrence of these disorders (8, 9). The exact underlying and interconnected genetic mechanisms remain under intense scientific study. However, some culprit genetic targets, as well as biological pathways have been identified and connected to the disorders’ occurrence, such as the serotonergic and dopaminergic system (10). Related studies have concentrated on a small number of genes in the dopamine and serotonin pathways, such as *COMT* Val158Met (11) and *DRD2* C957T (12), *DRD4* (13) (14), *HTR2A* (15), or the Val66Met polymorphism in *BDNF* (16). However, despite the extended research, findings have still been inconclusive. Both cognitive and behavioral disorders are multifactorial, and thus it becomes difficult to describe and sometimes distinguish them. These disorders can co-occur in a person, a fact that can be evidenced, not only by the common genes or single nucleotide polymorphisms (SNPs) that they share, but also through their related ontologies (**Figures 1 and 2**).

**Figure 1.**
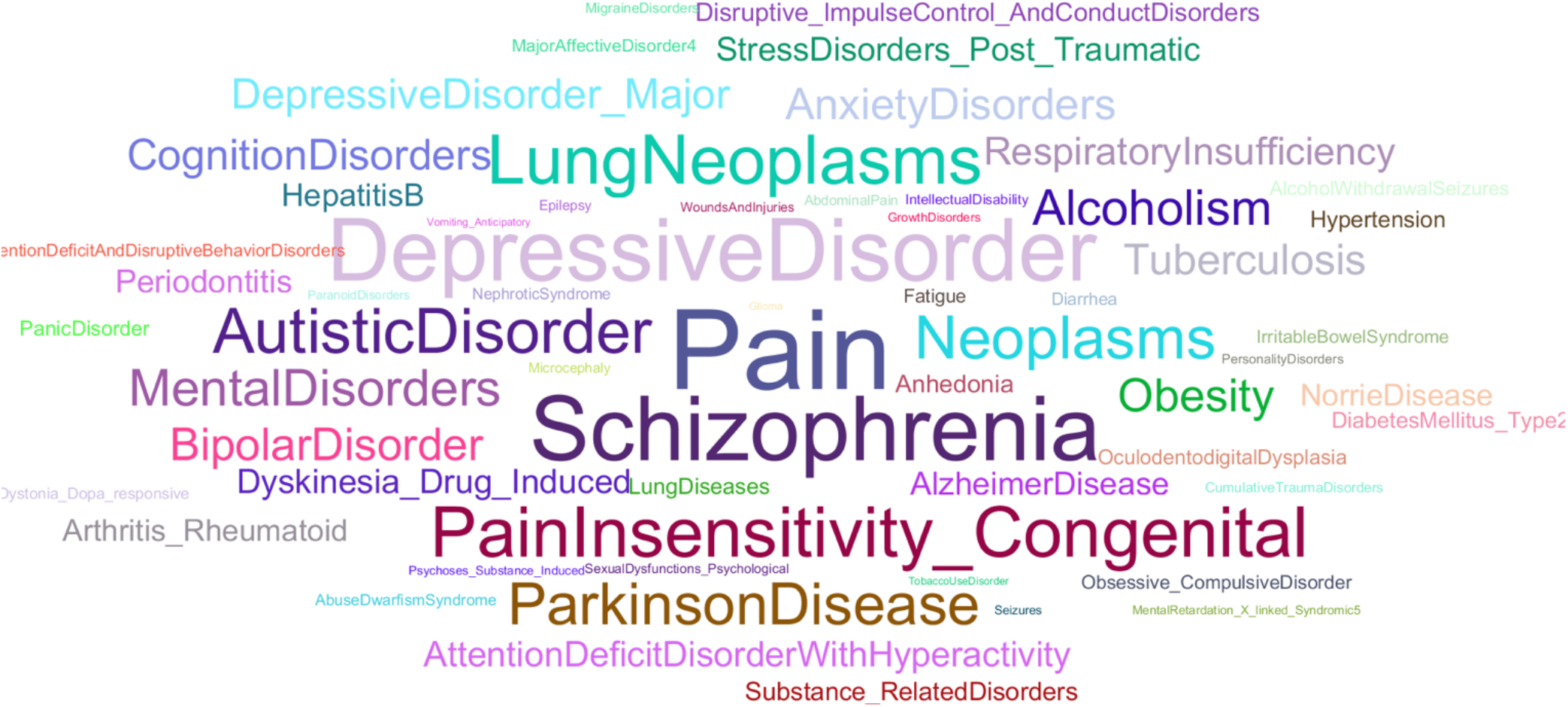
Ontologies which are connected to Behavioral Disorders, visualized in Watermark presentation graph.

**Figure 2.**
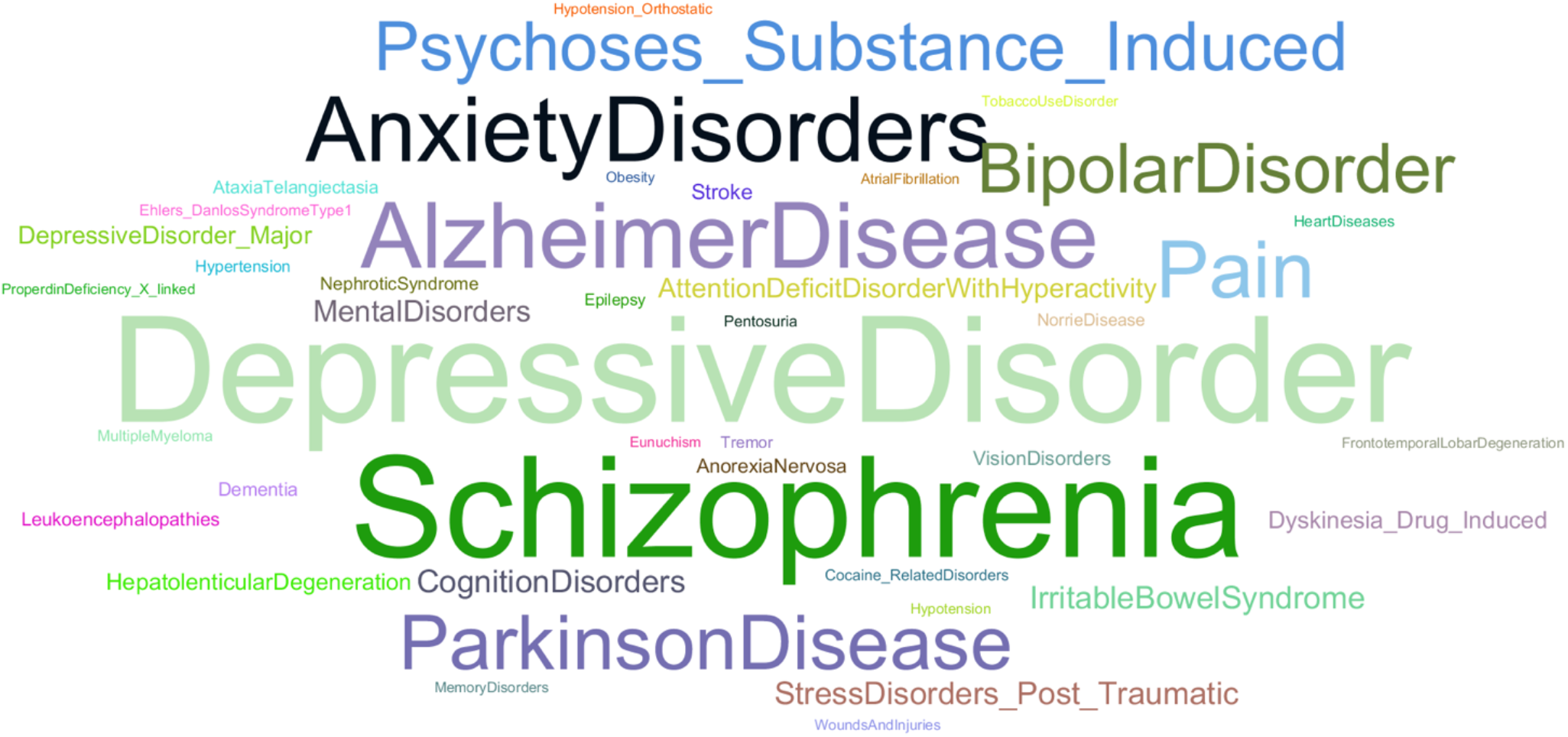
Ontologies which are connected to Cognitive Disorders, visualized in Watermark presentation graph.

Knowledge regarding genetics and genomics in the last two centuries has established a strong basis which allows the development of novel techniques and applications. In the context of research and clinical practice, the availability of modern biotechnological tools such as whole-genome sequencing (WGS) and whole-exome sequencing (WES) analysis, has helped scientists to better understand the molecular background and genetics of a disease, a feat which in turn enables the development of novel management and therapeutic approaches under the scope of personalized medicine (17). Another decisive factor towards this goal was the completion of the Human Genome project, which yielded massive amounts of genomic data and enriched the knowledge of genetic variants and their impact on human life and disease. As long as the accumulation of genomic data continues, the need to create and implement new methods, techniques and pipelines will grow stronger. Naturally, when exploring genetic variation and its potential effects, one cannot limit themselves to the coding regions of the human genome. As it has been evidenced, polymorphisms occurring in regions that do not code for proteins, such as promoters, splice sites, intergenic regions and introns, can have equally complex and strong effects (18-21). When analyzing genetic variation in the context of disease and disorder, the term “genomic grammar” can be appropriate, since it is not limited to the gene but encompasses factors related to the dark part of the genome. The “genomic grammar” of diseases and disorders, when combined with other disciplines and analyses, can be a powerful tool towards the emerging field of precision medicine. To navigate this ambiguous and complex genomic terrain we present a novel bioinformatic approach, through an integrated pipeline consisting of systemic data and text mining procedures, annotation progress and semantic analysis. The end goal of the strategy described herein is to goal to identify the most associated SNPs and their genetic variants, gaining knowledge which in the future can potentially support the clinical genomic diagnosis, and introduce new pharmacological targets for prevention and treatment of cognitive and behavioral disorders. The general scheme of the integrated bioinformatic approach is presented in **Figure 3**.

**Figure 3.**
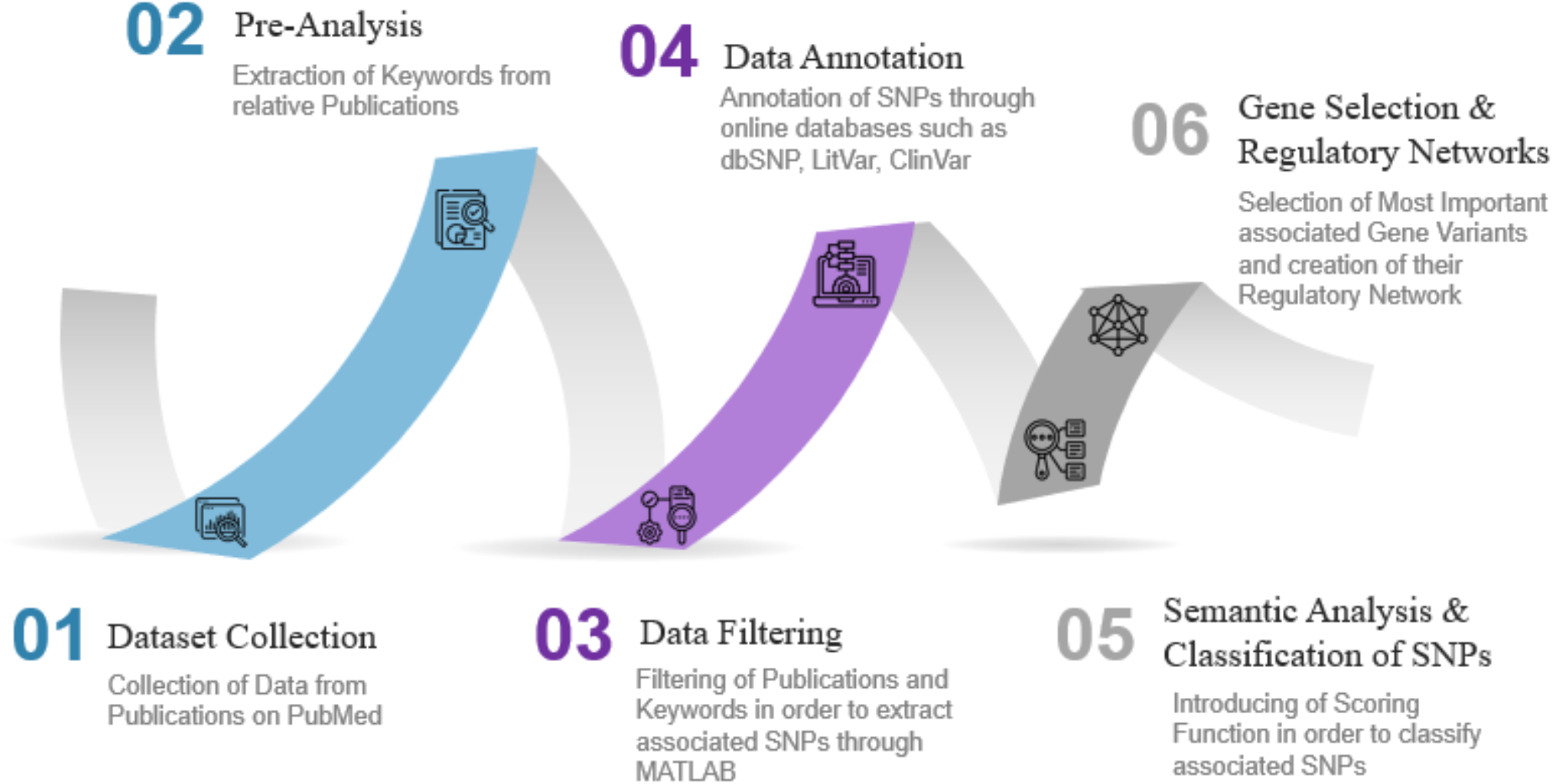
The method of the bioinformatic analysis, presented in 6 steps, displayed in a flow chart presentation.

## Methods

### Dataset Collection

The MEDLINE and PubMed databases were searched for English-language publications that contained the key terms “Behavioral Disorders”, Cognitive Disorders” with no date restriction. Publications associated with these terms were downloaded from online database PubMed (https://pubmed.ncbi.nlm.nih.gov/). PubMed, as a part of NCBI (National Center for Biotechnology Information), is a free access search engine which enables users to acquire admission to a database consisting of MEDLINE files, with summaries, reports or even entire articles on life sciences and biomedicine issues (22). This data was downloaded in MEDLINE format and stored into Excel sheets.

### Pre-Analysis

In this step, the extraction of keywords associated with cognitive and behavioral disorders was performed. The process took place in MATLAB Bioinformatics toolbox, a numeric computer environment through text mining analysis, from the associated publications collected from PubMed. MATLAB is also a computer programming language which uses algorithms and calculations in order to analyze large data volumes and visualize them into appealing formats (23). MATLAB was programmed to extract the keywords individually for each disorder. The most important and relevant keywords for each disorder were represented in Bubble Chart diagrams, using Tableau Software interactive data visualization software. Common keywords were also recorded, identified and collected into a table. MATLAB is also a computer programming language which uses algorithms and calculations in order to analyze large data volumes and visualize them into appealing formats (23).

### Data Filtering

The available studies’ abstracts were filtered specifically for human related studies and were edited through data mining and semantic methods in MATLAB, in order to identify those that refer to genes by using a dictionary of the gene, allele and pseudogene names for *Homo Sapiens* from the Gene database of the National Center for Biotechnology Information (NCBI), and also those that contained SNP variants. The identified genes, SNPs and variants were extracted through a targeted query search by text mining analysis, using regular expressions by combining each gene or variant with their synonyms and the keywords “Behavioral” and/or “Cognitive”. A second-level analysis was performed in order to estimate the internal links between genes, through selected publications. Internal links were created when genes, alleles, pseudogenes, or transcription factors were mentioned in the same publication. Those genetic variants were stored into sheet datasets in Excel.

### Data Annotation

Furthermore, the extracted genes, SNPs and variants were mined for extraction of additional information from available online databases, such as dbSNP, GWAS-Catalog, LitVar and ClinVar (22, 24-26). The additional information contained the “snp_name”, “frequency”, “chromosome”, “position”, “reference genome”, “change”, “gene_name”, “variant_type”, “disease”, “litvar”, the existence or not of corresponding clinical data from ClinVar database as “ClinVar”. The final dataset of SNPs and variants associated with cognitive and behavioral disorders was annotated through a MATLAB algorithm which connected them with the additional information from the online databases.

### Semantic Analysis and Classification of SNPs

In this step, the annotated SNPs and variants were scored through semantic analysis in order to identify and categorize the most “suspected” genetic targets associated with the occurrence of cognitive and behavioral disorders. Thus, a scoring function was built which is described below. Through this scoring, the identified classes of SNPs (“Strong-associated SNPs”, “High-associated SNPs” and “Associated SNPs”) have been structured and evaluated for their clinical severity.

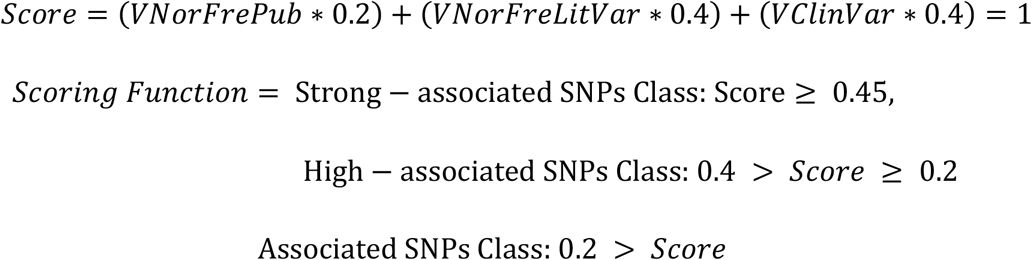

where:

V*NorFrePub* = The Normalized frequency of the identified SNPs from the PubMed dataset (Max=1, Min=0), *VNorFreLitVar* = The Normalized frequency of the identified SNPs that are linked to Cognitive and Behavioral Disorders from the LitVar Database (Scalar value, Max=1, Min=0), *VClinVar* = Boolean Parameter, 1 = The SNP was identified in the ClinVar databases and is connected to Behavioral or/and Cognitive Disorder, 0 = no connection to the ClinVar, or no connection to Behavioral or/and Cognitive Disorder, -1=The SNP was identified in the ClinVar databases and is not connected to Behavioral or/and Cognitive Disorders

All the identified SNPs were classified in these three major classes based on the annotated information contained into the Excel sheet datasets. Although the SNPs connected to each disorder are unique, our analysis identified genetic variants which participate in both disorders, a fact which makes the construction of the respective “genomic grammars” challenging. This genetic overlap is an indication that these disorders are somehow connected to each other, not only through their ontologies, but also through their genetic basis.

### Gene Selection and Regulatory Networks

Each SNP variant corresponds to a gene variant, based on the annotation progress through the online database dbSNP. These genetic targets, combined with the mined gene variants extracted from the MATLAB algorithm, have been grouped according to each disorder. Via classification process and “strongly-associated SNP” variants, the strongly suspected genetic variants were identified and collected into lists. For the regulatory networks of these genetic targets to be built, two searches were performed on the website GeneMANIA, individually for each disorder. GeneMANIA is a user-friendly website that allows the user to create a complex network of gene interactions from an existing gene list. Through this creation of the network, the user can assume various gene functions, analyze gene catalogs but also classify genes, hierarchically, according to their functions. The resulting network includes genes, which are more related to each other, with the initial list which has been submitted, accompanied by detailed information from a variety of genomic and proteomic databases (27).

## Results

### Dataset Collection

Through systematic data mining and semantic analysis techniques, the most frequently reported gene variants and SNPs of cognitive and behavioral disorders were identified. On the basis of containing the terms “behavioral disorders” or “cognitive disorders” in the title or abstract of the MEDLINE files, a total number of 1,359,425 and 199,267 publications were retrieved on behavioral and cognitive disorders respectively

### Pre-Analysis

A total number of 5551 keywords were found which are directly connected to cognitive disorders, as well as 7660 keywords to behavioral. In **Table 1**, the most frequently shown key terms, describing either cognitive or behavioral disorders within the extracted dataset, are shown. The same key terms were visualized into Bubble Charts through Tableau software (**Figures 4**,**5**). Furthermore, common keywords that applied to both disorders were also identified and recorded into **Table 2**.

**Table 1.**
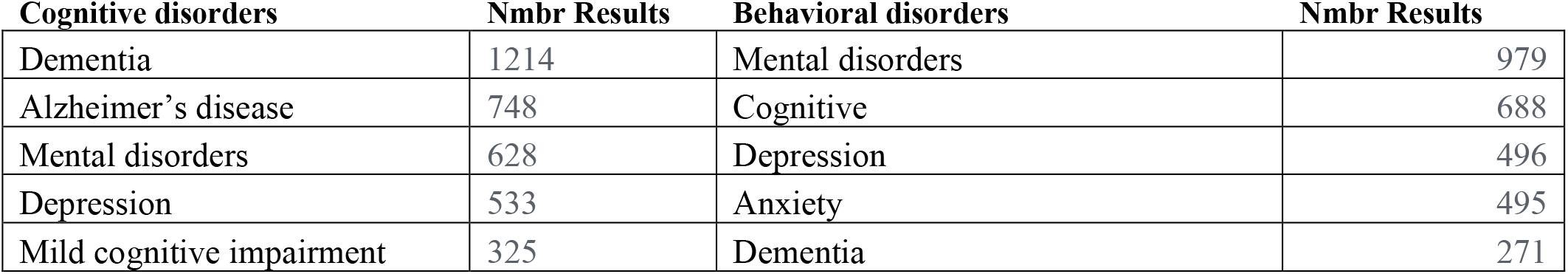
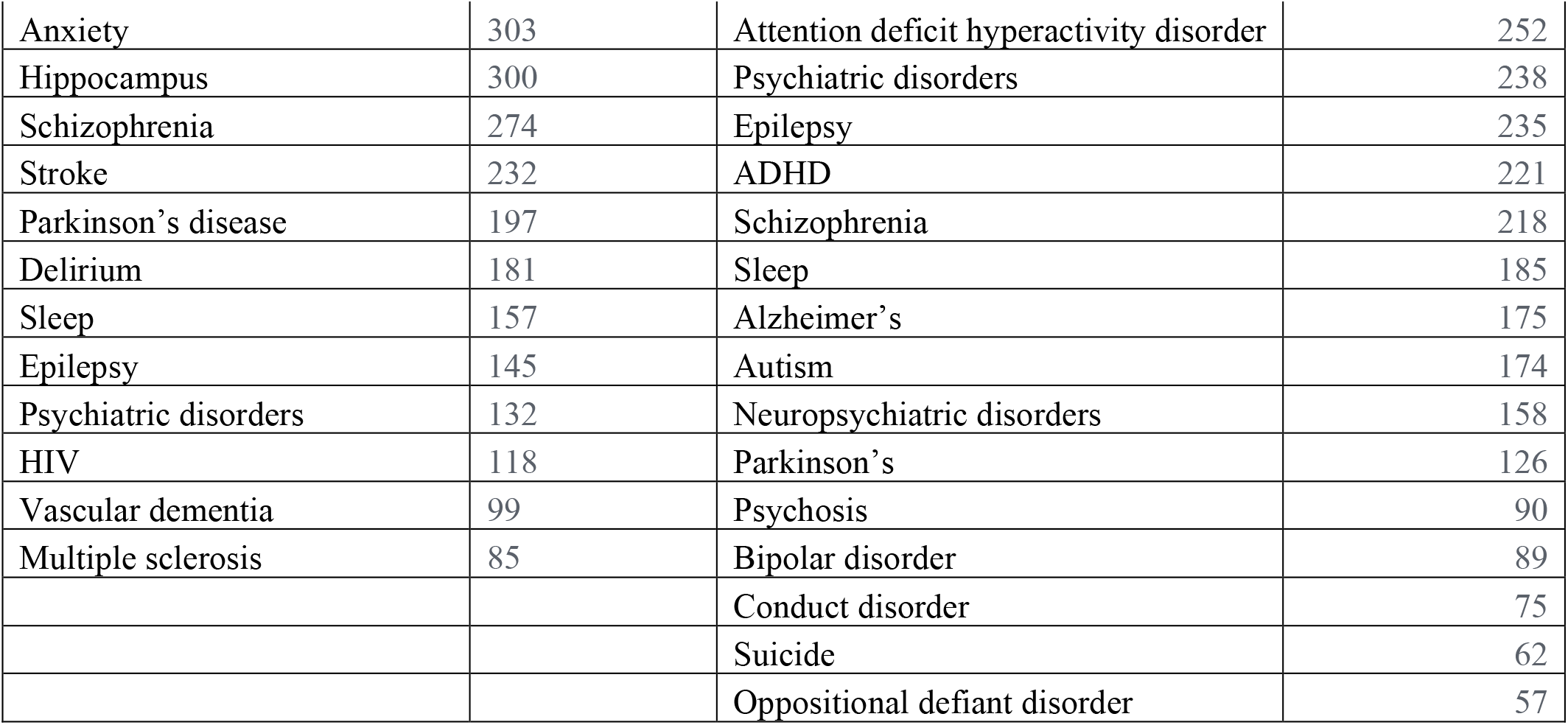
List of the most frequently shown key terms describing cognitive and behavioral disorders within the dataset.

| **Cognitive disorders** | **Nmbr Results** | **Behavioral disorders** | **Nmbr Results** |
| --- | --- | --- | --- |
| Dementia | 1214 | Mental disorders | 979 |
| Alzheimer’s disease | 748 | Cognitive | 688 |
| Mental disorders | 628 | Depression | 496 |
| Depression | 533 | Anxiety | 495 |
| Mild cognitive impairment | 325 | Dementia | 271 |
| Anxiety | 303 | Attention deficit hyperactivity disorder | 252 |
| Hippocampus | 300 | Psychiatric disorders | 238 |
| Schizophrenia | 274 | Epilepsy | 235 |
| Stroke | 232 | ADHD | 221 |
| Parkinson's disease | 197 | Schizophrenia | 218 |
| Delirium | 181 | Sleep | 185 |
| Sleep | 157 | Alzheimer's | 175 |
| Epilepsy | 145 | Autism | 174 |
| Psychiatric disorders | 132 | Neuropsychiatric disorders | 158 |
| HIV | 118 | Parkinson's | 126 |
| Vascular dementia | 99 | Psychosis | 90 |
| Multiple sclerosis | 85 | Bipolar disorder | 89 |
|  |  | Conduct disorder | 75 |
|  |  | Suicide | 62 |
|  |  | Oppositional defiant disorder | 57 |

**Figure 4.**
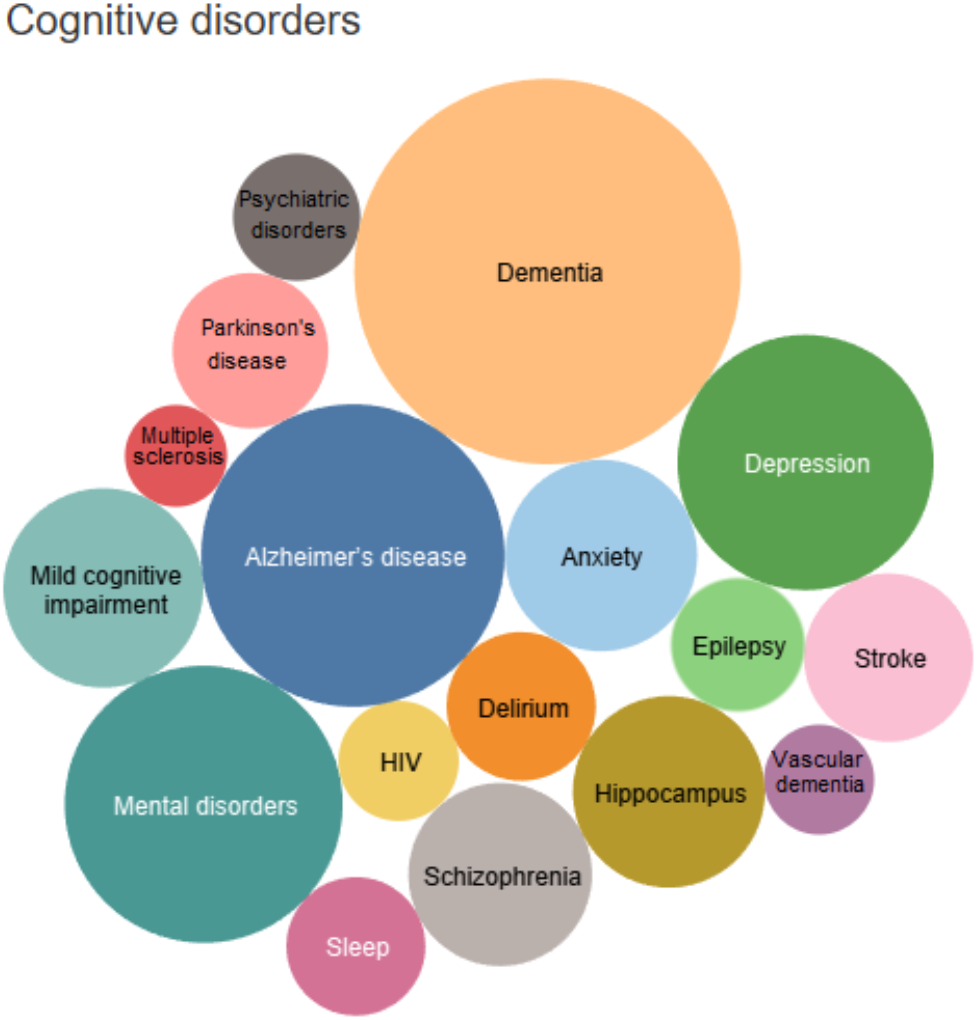
The key terms that are mostly associated with Cognitive Disorders, in Bubble Chart form. The size of the circles represents the intensity of the corresponding term’s frequency.

**Figure 5.**
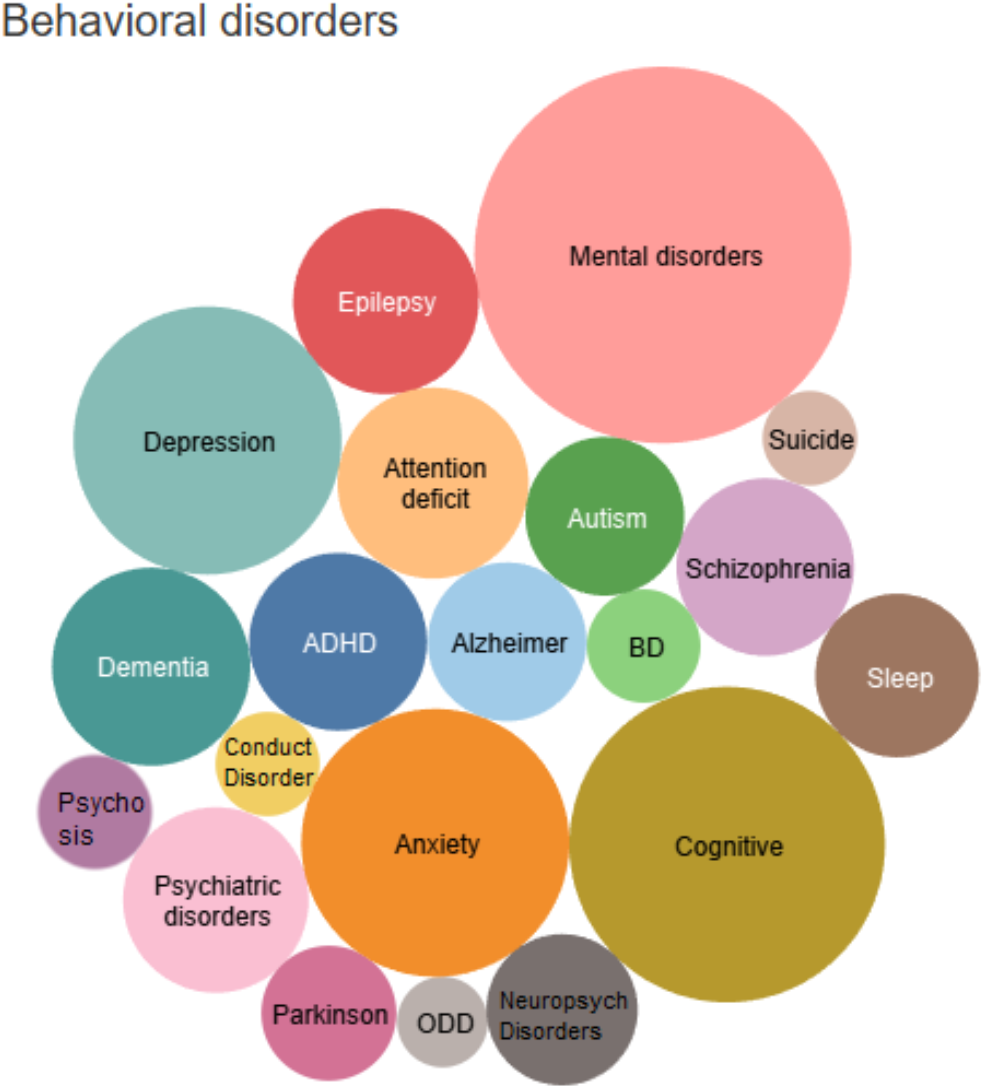
The key terms that are mostly associated with Behavioral Disorders, in Bubble Chart form. The size of the circles represents the intensity of the corresponding term’s frequency.

**Table 2.**
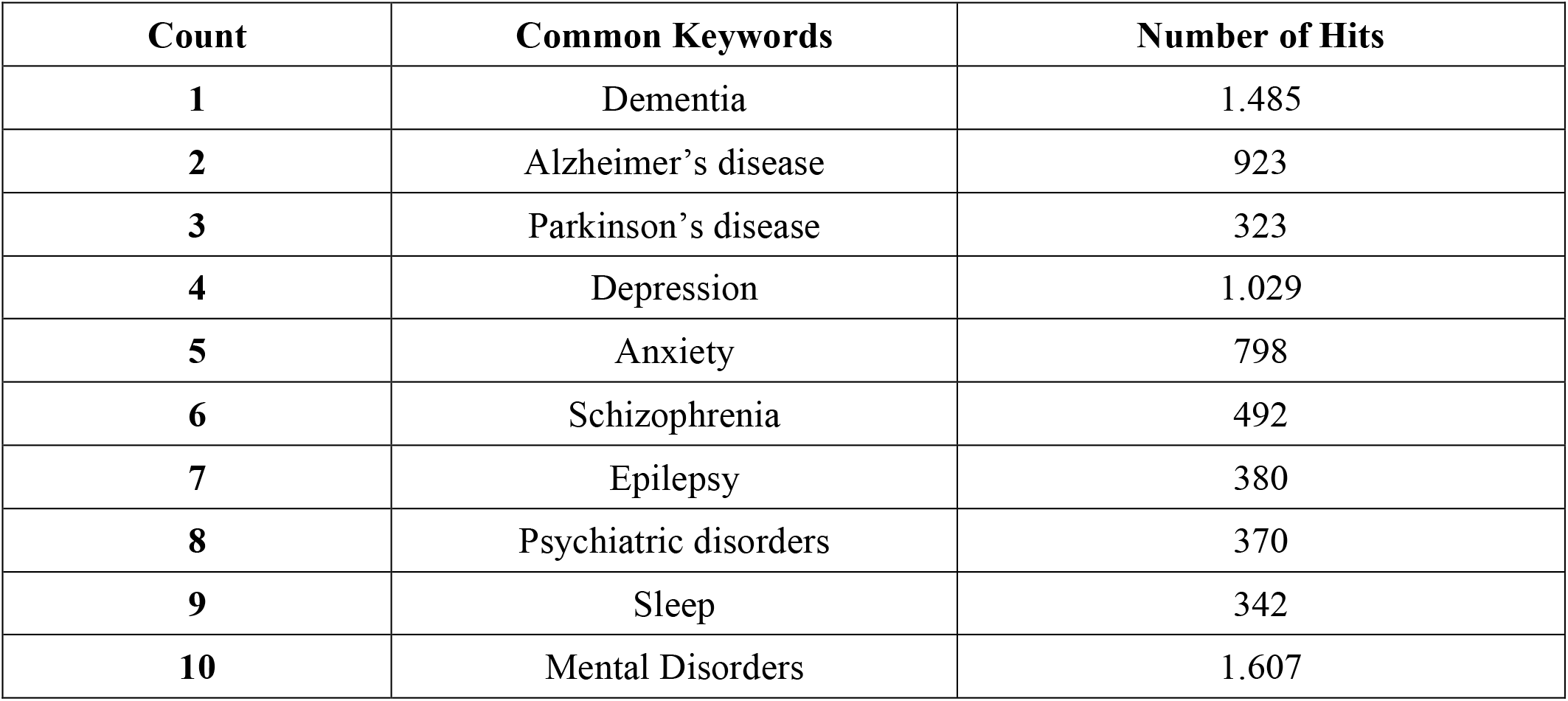
List of common key terms describing both cognitive and behavioral disorders.

| Count | Common Keywords | Number of Hits |
| --- | --- | --- |
| 1 | Dementia | 1.485 |
| 2 | Alzheimer's disease | 923 |
| 3 | Parkinson's disease | 323 |
| 4 | Depression | 1.029 |
| 5 | Anxiety | 798 |
| 6 | Schizophrenia | 492 |
| 7 | Epilepsy | 380 |
| 8 | Psychiatric disorders | 370 |
| 9 | Sleep | 342 |
| 10 | Mental Disorders | 1.607 |

### Data Filtering

Through filtering of the dataset collection, 82 SNPs and 1100 genes, alleles, pseudogenes and transcription factors associated with cognitive and behavioral disorders were reported and imported from online databases. More specifically, 30 cognitive disorder-associated and 52 behavioral disorder-associated SNPs have been identified. Furthermore, 502 cognitive disorder-associated genetic variants and 598 behavioral disorder-associated genetic variants were identified. The results of SNPs concerning cognitive disorders are shown into **Table 3**, whereas the results concerning behavioral disorders into **Table 4**.

**Table 3.** List of SNPs mostly connected to Cognitive disorders, accompanied with the frequency of their appearance within the dataset.

| SNPs | Frequency |
| --- | --- |
| rs25531 | 7 |
| rs16917204 | 6 |
| rs11030104 | 6 |
| rs16917237 | 6 |
| rs2049045 | 6 |
| rs6265 | 6 |
| rs988748 | 6 |
| rs6484320 | 6 |
| rs7103411 | 6 |
| rs7282236 | 5 |
| rs9675262 | 5 |
| rs9820240 | 5 |
| rs10916409 | 5 |
| rs540422 | 5 |
| rs7220676 | 5 |
| rs2968869 | 4 |
| rs1334071 | 4 |
| rs16933774 | 4 |
| rs10932201 | 4 |
| rs2551645 | 4 |
| rs2254137 | 4 |
| rs6740584 | 4 |
| rs2551640 | 4 |
| rs11136000 | 3 |
| rs28363170 | 3 |
| rs3836790 | 3 |
| rs921451 | 3 |
| rs3837091 | 3 |
| rs1799836 | 3 |
| rs4680 | 3 |

**Table 4.** List of SNPs mostly connected to Behavioral disorders, accompanied with the frequency of their appearance within the dataset

| SNPs | Frequency |
| --- | --- |
| rs2144025 | 5 |
| rs2104425 | 5 |
| rs1046706 | 4 |
| rs2360867 | 4 |
| rs4140535 | 4 |
| rs1778258 | 4 |
| rs17273700 | 4 |
| rs1228814 | 4 |
| rs11568817 | 4 |
| rs130058 | 4 |
| rs3746554 | 3 |
| rs1051312 | 3 |
| rs179991 | 3 |
| rs2268498 | 2 |
| rs1801028 | 2 |
| rs2268498 | 2 |
| rs4537731 | 2 |
| rs21105 | 2 |
| rs684302 | 2 |
| rs1800532 | 2 |
| rs1535255 | 2 |
| rs2023239 | 2 |
| rs1049353 | 2 |
| rs806368 | 2 |
| rs1386494 | 2 |
| rs2430561 | 2 |
| rs334558 | 2 |
| rs4532 | 2 |
| rs4867798 | 2 |
| rs265981 | 2 |
| rs1800497 | 2 |
| rs104894220 | 2 |
| rs144999500 | 2 |
| rs3732783 | 2 |
| rs6280 | 2 |
| rs1800443 | 2 |
| rs144132215 | 2 |
| rs7301328 | 2 |
| rs53576 | 2 |
| rs7118422 | 1 |
| rs16969968 | 1 |
| rs4713902 | 1 |
| rs1799971 | 1 |
| rs2402959 | 1 |
| rs6965643 | 1 |
| rs2295633 | 1 |
| rs45573936 | 1 |
| rs10508649 | 1 |
| rs2491397 | 1 |
| rs3734967 | 1 |
| rs3813034 | 1 |
| rs1176744 | 1 |

### Data Annotation

The identified and filtered SNPs associated with cognitive disorders were annotated via MATLAB algorithms which imported more detailed information for them from online databases. The process was accordingly repeated for behavioral disorders. In **Tables 5**,**6** the annotated SNPs from dbSNP, LitVar and ClinVar, and their genetic variant annotation, are shown. The annotated SNPs are also displayed, graphically in Pie Charts, in **Figures 6** and **7**, for behavioral and cognitive disorders, respectively. A result of great interest is the distribution of the extracted SNPs, connected to these disorders, among the human chromosomes, which is demonstrated in **Figure 8**.

**Table 5.** List of the annotated SNPs, associated with Behavioral disorders, from online databases, accompanied with their scoring function.

| SNPs | W1 | W2 | W3 | Scoring Function | Genes |
| --- | --- | --- | --- | --- | --- |
| rs1800497 | 0,25 | 1 | 1 | 0,85 | ANKK1 |
| rs6280 | 0,25 | 1 | 1 | 0,85 | DRD3 |
| rs1801028 | 0,25 | 1 | 1 | 0,85 | DRD2 |
| rs1799971 | 0 | 1 | 1 | 0,8 | OPRM1 |
| rs7301328 | 0,25 | 0,5 | 1 | 0,65 | GRIN2B |
| rs130058 | 0,75 | 0,75 | 0 | 0,45 | HTR1B |
| rs53576 | 0,25 | 1 | 0 | 0,45 | OXTR |
| rs1800532 | 0,25 | 1 | 0 | 0,45 | TPH1 |
| rs334558 | 0,25 | 1 | 0 | 0,45 | GSK3B |
| rs1049353 | 0,25 | 1 | 0 | 0,45 | CNR1 |
| rs3732783 | 0,25 | 0 | 1 | 0,45 | DRD3 |
| rs1800443 | 0,25 | 0 | 1 | 0,45 | DRD4 |
| rs2144025 | 1 | 0,5 | 0 | 0,4 | ESR1 |
| rs2491397 | 0 | 0 | 1 | 0,4 | GABBR2 |
| rs3734967 | 0 | 0 | 1 | 0,4 | HTR5A-AS1 |
| rs11568817 | 0,75 | 0,5 | 0 | 0,35 | HTR1B |
| rs2023239 | 0,25 | 0,75 | 0 | 0,35 | CNR1 |
| rs806368 | 0,25 | 0,75 | 0 | 0,35 | CNR1 |
| rs1051312 | 0,5 | 0,5 | 0 | 0,3 | SNAP25 |
| rs1176744 | 0 | 0,75 | 0 | 0,3 | HTR3B |
| rs3813034 | 0 | 0,75 | 0 | 0,3 | SLC6A4 |
| rs1046706 | 0,75 | 0,25 | 0 | 0,25 | SLC9A9 |
| rs2360867 | 0,75 | 0,25 | 0 | 0,25 | SLC9A9 |
| rs1386494 | 0,25 | 0,5 | 0 | 0,25 | TPH2 |
| rs1535255 | 0,25 | 0,5 | 0 | 0,25 | CNR1 |
| rs2268498 | 0,25 | 0,5 | 0 | 0,25 | OXTR |
| rs2104425 | 1 | 0 | 0 | 0,2 | LOC105375050 |
| rs1228814 | 0,75 | 0 | 0 | 0,15 | HTR1B |
| rs1778258 | 0,75 | 0 | 0 | 0,15 | HTR1B |
| rs4140535 | 0,75 | 0 | 0 | 0,15 | HTR1B |
| rs17273700 | 0,75 | 0 | 0 | 0,15 | HTR1B |
| rs4867798 | 0,25 | 0,25 | 0 | 0,15 | DRD1 |
| rs2268498 | 0,25 | 0,25 | 0 | 0,15 | OXTR |
| rs265981 | 0,25 | 0,25 | 0 | 0,15 | DRD1 |
| rs684302 | 0,25 | 0,25 | 0 | 0,15 | TPH1 |
| rs3746554 | 0,5 | 0 | 0 | 0,1 | ZNFX1 |
| rs179991 | 0,5 | 0 | 0 | 0,1 | ATXN1 |
| rs2295633 | 0 | 0,25 | 0 | 0,1 | FAAH |
| rs4537731 | 0,25 | 0 | 0 | 0,05 | TPH1 |
| rs144999500 | 0,25 | 0 | 0 | 0,05 | DRD2 |
| rs144132215 | 0,25 | 0 | 0 | 0,05 | DRD5 |
| rs21105 | 0,25 | 0 | 0 | 0,05 | N/A |
| rs2430561 | 0,25 | 0 | 0 | 0,05 | IFNG |
| rs4532 | 0,25 | 1 | -1 | 0,05 | DRD1 |
| rs4713902 | 0 | 0 | 0 | 0 | FKBP5 |
| rs45573936 | 0 | 0 | 0 | 0 | SLC29A1 |
| rs2402959 | 0 | 0 | 0 | 0 | LOC105375501 |
| rs6965643 | 0 | 0 | 0 | 0 | LOC105375501 |
| rs7118422 | 0 | 0 | 0 | 0 | STIM1 |
| rs10508649 | 0 | 0 | 0 | 0 | PIP4K2A |
| rs16969968 | 0 | 0,5 | -1 | -0,2 | CHRNA5 |
| rs104894220 | 0,25 | 0 | -1 | -0,35 | DRD2 |

**Table 6.** List of the annotated SNPs, associated with Cognitive disorders, from online databases, accompanied with their scoring function.

| **SNPs** | **W1** | **W2** | **W3** | **Scoring**<br>**Function** | **Genes** |
| --- | --- | --- | --- | --- | --- |
| rs6265 | 1 | 1 | 1 | 1 | BDNF |
| rs4680 | 0,25 | 1 | 1 | 0,85 | COMT |
| rs25531 | 1 | 1 | 0 | 0,6 | SLC6A4 |
| rs11136000 | 0,25 | 1 | 0 | 0,45 | CLU |
| rs16917204 | 1 | 0,5 | 0 | 0,4 | BDNF-AS |
| rs7103411 | 1 | 0,5 | 0 | 0,4 | BDNF |
| rs16917237 | 1 | 0,5 | 0 | 0,4 | BDNF |
| rs1799836 | 0,25 | 0,75 | 0 | 0,35 | MAOB |
| rs11030104 | 1 | 0,25 | 0 | 0,3 | BDNF |
| rs988748 | 1 | 0,25 | 0 | 0,3 | BDNF |
| rs28363170 | 0,25 | 0,5 | 0 | 0,25 | SLC6A3 |
| rs7220676 | 0,75 | 0,25 | 0 | 0,25 | N/A |
| rs6740584 | 0,5 | 0,25 | 0 | 0,2 | CREB1 |
| rs2049045 | 1 | 0 | 0 | 0,2 | BDNF |
| rs6484320 | 1 | 0 | 0 | 0,2 | BDNF |
| rs3836790 | 0,25 | 0,25 | 0 | 0,15 | SLC6A3 |
| rs540422 | 0,75 | 0 | 0 | 0,15 | PICALM |
| rs9675262 | 0,75 | 0 | 0 | 0,15 | N/A |
| rs7282236 | 0,75 | 0 | 0 | 0,15 | N/A |
| rs10916409 | 0,75 | 0 | 0 | 0,15 | N/A |
| rs9820240 | 0,75 | 0 | 0 | 0,15 | DCLK3 |
| rs2968869 | 0,5 | 0 | 0 | 0,1 | ERC1 |
| rs2254137 | 0,5 | 0 | 0 | 0,1 | CREB1 |
| rs10932201 | 0,5 | 0 | 0 | 0,1 | CREB1 |
| rs2551645 | 0,5 | 0 | 0 | 0,1 | CREB1 |
| rs2551640 | 0,5 | 0 | 0 | 0,1 | CREB1 |
| rs16933774 | 0,5 | 0 | 0 | 0,1 | ASTN2 |
| rs1334071 | 0,5 | 0 | 0 | 0,1 | ASTN2 |
| rs921451 | 0,25 | 0 | 0 | 0,05 | DDC |
| rs3837091 | 0,25 | 0,5 | -1 | -0,15 | DDC |

**Figure 6.**
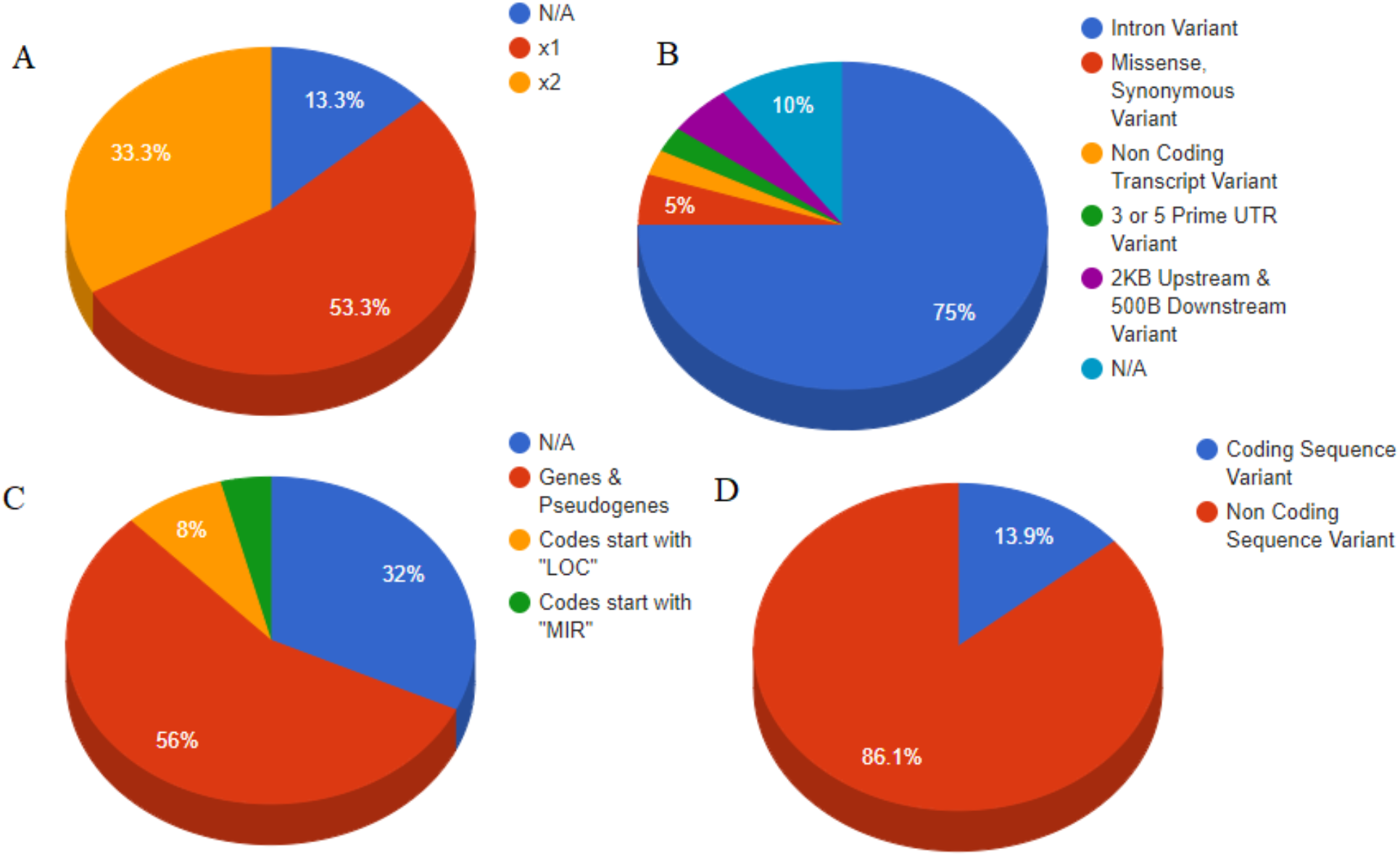
Database analysis results for cognitive disorders. (A)”x1”, “x2” corresponds to the number of the affected regions per SNP. (B) The six identified categories of SNPs (C) The types of SNP categories (D) SNPs analysis per two major genomic regions.

**Figure 7.**
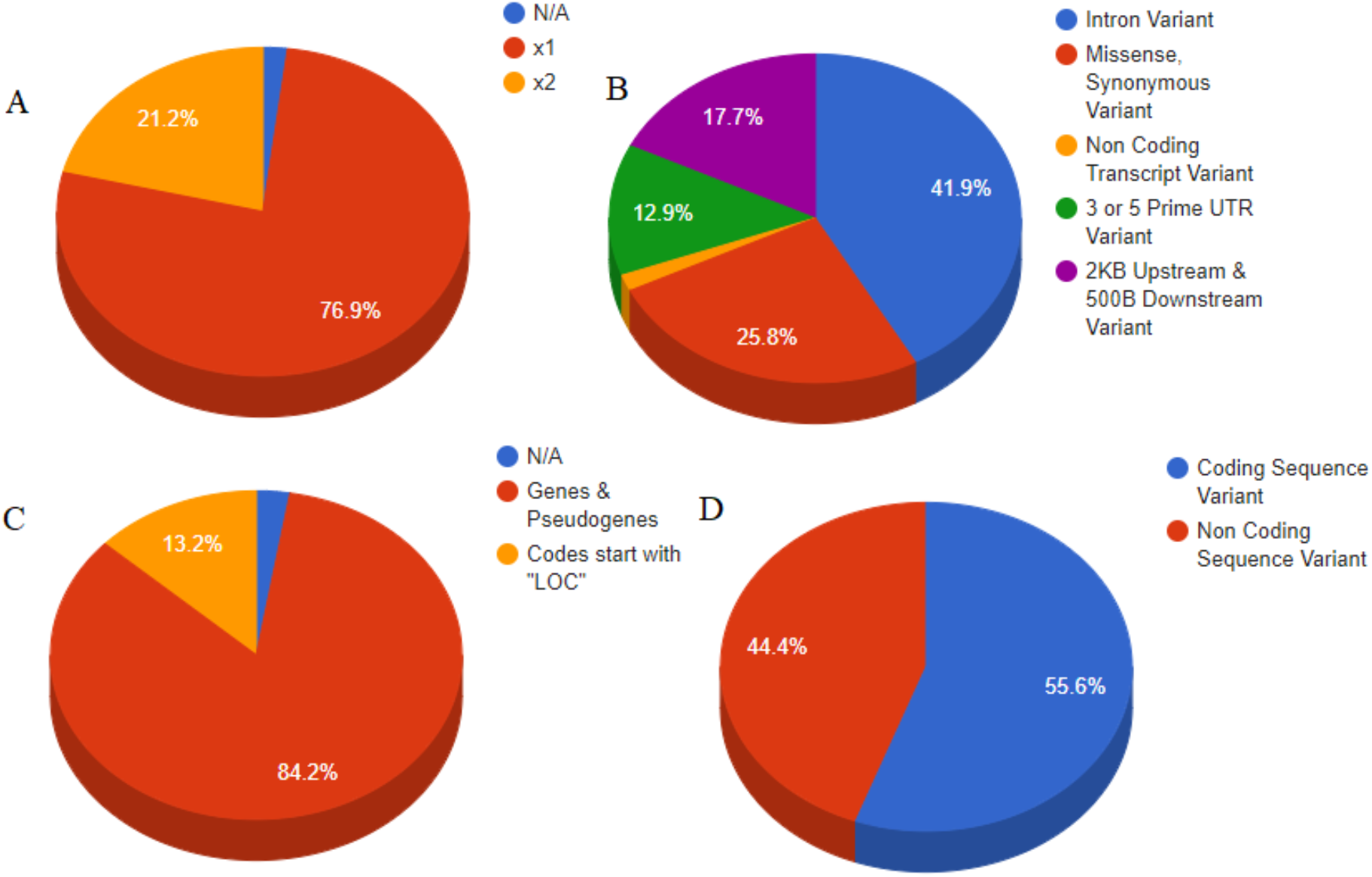
Database analysis results for behavioral disorders. (A)”x1”, “x2” corresponds to the number of the affected regions per SNP. (B) The five identified categories of SNPs (C) The types of SNP categories (D) SNPs analysis per two major genomic regions.

**Figure 8.**
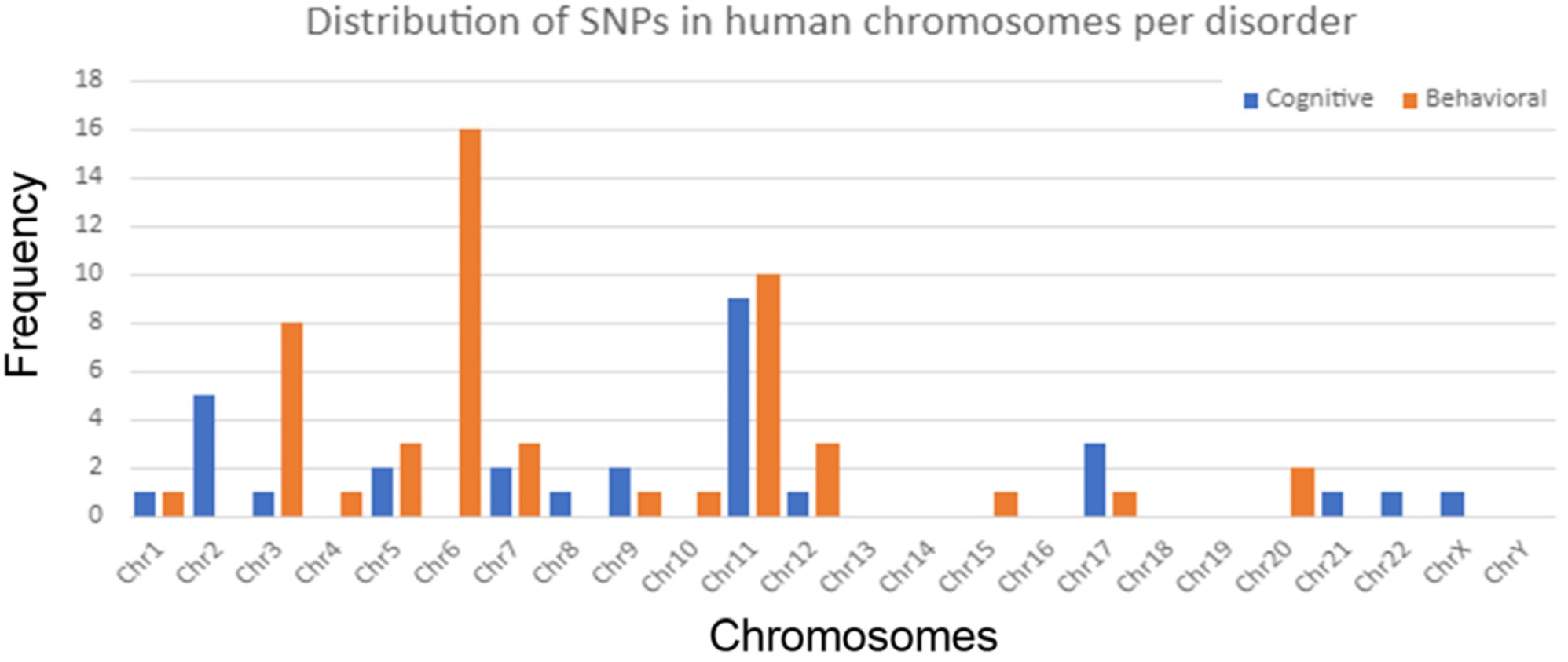
Distribution of SNPs connected to cognitive and behavioral disorders throughout human chromosomes.

### Semantic Analysis and Classification of SNPs

The implemented scoring function allowed the selection and distinction of the most “suspicious” SNP variants, connected to cognitive and behavioral disorders. The scores of the SNP variants in both disorders are also shown in **Tables 5** and **6**. Classification of the annotated SNP variants was formed into three major classes: “highly-associated SNPs”, “most-associated SNPs” and “associated SNPs”, based on their variant scores. Only 7 SNP variants (23,33%) were found to be highly associated with cognitive disorders, in addition to 14 SNP variants (27%) that were highly associated with behavioral disorders. In **Figure 9**, the results of the classification are visualized into Pie Charts.

**Figure 9.**
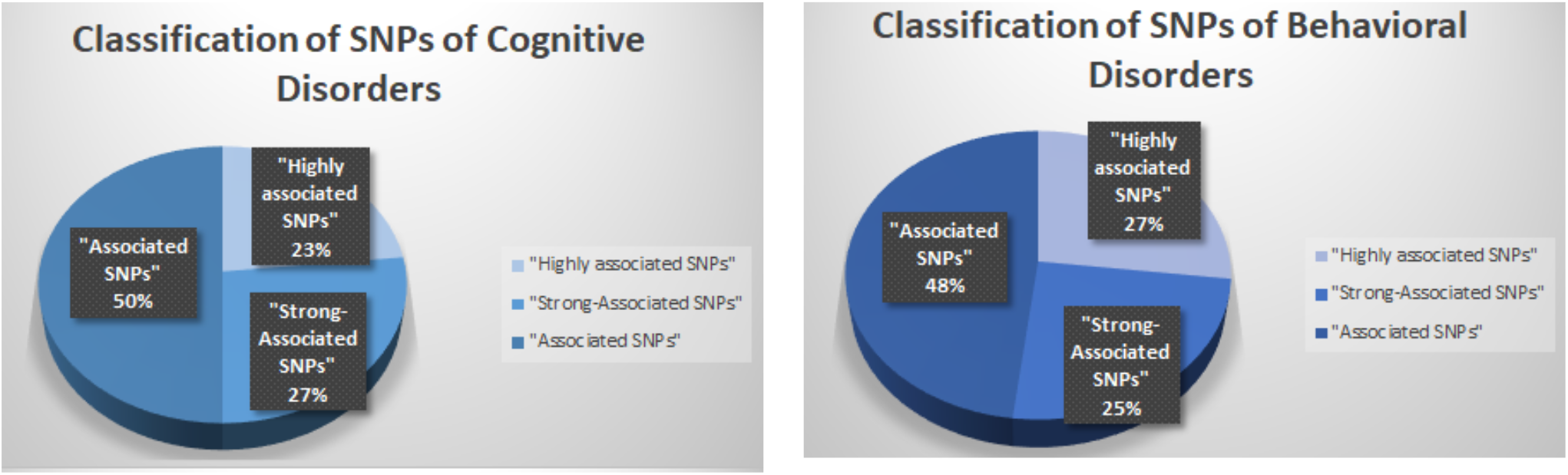
Classification of SNPs associated with Cognitive and Behavioral Disorders in three major categories: “highly-associated”, “strong-associated” and “associated”, presented in two Pie Charts.

### Gene Selection and Regulatory Networks

Through identification and classification of most-associated SNPs with the disorders we study, we were allowed to also identify the most strongly “suspected” genetic variants that could potentially predispose to cognitive and/or behavioral disorders. In our study, the genetic variants *APOE, BDNF, COMT* and *SLC6A4* were selected as most connected to cognitive disorders, whereas Dopamine Receptors D (DRDs), *ANKK1, MET* and *HTR1B* were found to be mostly connected to behavioral disorders. By calculating the regulatory networks of the genetic variants of each disorder, it was revealed that DRDs and BDNF were evidently connected to each other and have also shown the strongest genetic links to cognitive and behavioral disorders amongst the others, thus their regulatory network was calculated in a graph representation (**Figure 10**).

**Figure 10.**
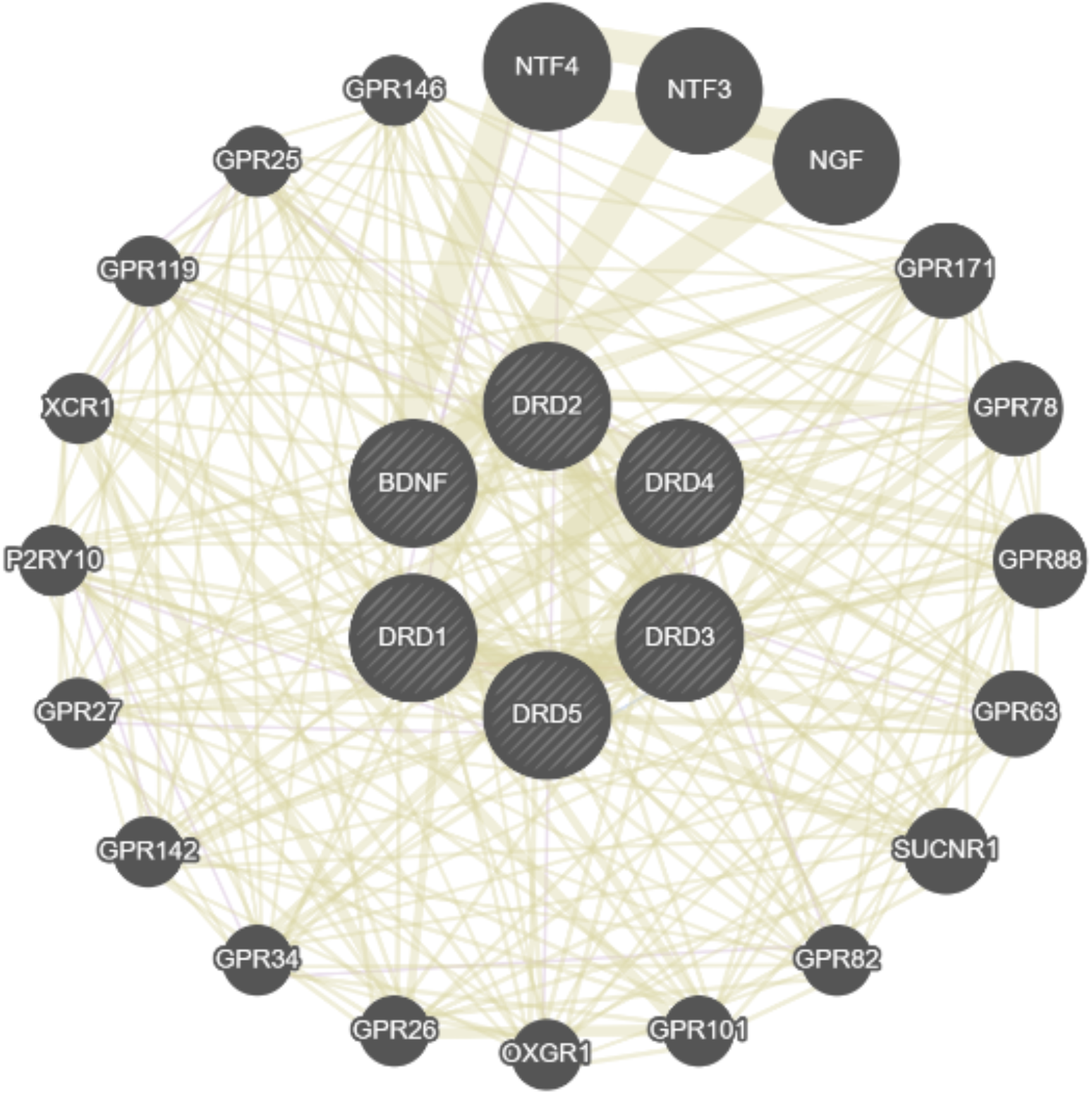
The regulatory network of the selected genetic variants strongly connected to cognitive and behavioral disorders, Dopamine Receptors D (DRD1, DRD2, DRD3, DRD4 and DRD5) and Brain Derived Neurotrophic Factor (BDNF), in a graph presentation.

## Discussion

The aim of this study was to explore the complex web of genetic causes underlying the occurrence of behavioral and cognitive disorders, a process which in the future can aid the proposition of novel pharmacological targets and the construction of innovative management and treatment strategies. Cognitive and behavioral disorders can potentially make their appearance due to a multifaceted genetic and epigenetic interplay. Tracking down the genetic and epigenetic causes that contribute to the display of such disorders is crucial in understanding and elucidating the various pathological conditions that may exist. Since a disorder, or generally, a disease can manifest as a result of a combination of adverse genetic polymorphisms, their identification and classification are significant for the efficient interpretation of the patient’s genomic profile in the clinical setting. In this study, we presented a new pipeline for the collection, filtering annotation, and evaluation of the candidate genetic targets towards the estimation of the significance of each genetic factor that is connected to the given disorders.

Initially, through the extraction of the most related key terms, an overlap between them was revealed, thanks to their ontologies. In **Figure 1** we can observe that the most connected key terms to behavioral disorders are “Pain”, “Schizophrenia” and “Depressive Disorders”, whereas in **Figure 2** the most associated key terms that apply to cognitive disorders are “Depressive Disorders”, “Schizophrenia” and “Anxiety Disorders”. As a fact, these disorders constitute parts of a greater set of mental disorders, which is confirmed in behavioral disorders from the key term “Pain”, that applies either to mental or physical pain. Of great interest are certain key terms extracted from cognitive disorders which subsequently appeared to be connected with behavioral ones, like “Depressive Disorders” and “Anxiety Disorders”, a fact which potentially indicates, inter alia, the existence of interplay between these disorders.

The second phase of this study, after the collection and annotation of the most strongly suspected genetic factors connected to these disorders, yielded interesting information. Of particular interest were the increased rates of SNPs found in non-coding areas of the genome (Cognitive Disorders: 86,1%, Behavioral Disorders: 44,4%). To date, we know that the non-coding regions of DNA make up 98% of the genome, as exome sequencing only examines less than 2% of the genome sequence and the current diagnostic contribution stands at about 30% (28). This region was originally called “junk DNA” as scientists considered this part of the genome useless. After a lot of studies on this part of DNA, the name of “junk DNA” has been switched to “dark DNA”, as, although the information we have about it is limited, which makes it rather unknown, it does not seem to be useless at all (29). In recent years, however, new studies have begun to identify more and more genetic elements or SNPs that belong to non-coding regions of DNA and are associated with a disease. In fact, it has been found that the majority of disease-related genetic polymorphisms, also identified by GWAS studies throughout the genome, are outside the regions that encode proteins (30). A huge amount of work is now being done to clarify all the functions of the genome, especially in the light of its polymorphisms within it, which may also be the causes of a disease or disorder. Ultimately, the “dark DNA”, and its association with the occurrence of diseases now becomes worthy of further investigation.

Additionally, the extracted SNPs were annotated for their position in human chromosomes. In **Figure 8**, we can observe that most of the SNPs related to cognitive disorders are found in chromosomes 11 and 2, whereas SNPs related to behavioral disorders are found in chromosomes 6, 11 and 3. As previously evidenced by studies, chromosome 6 has been linked to the onset of bipolar disorder, schizophrenia and depression (31), of alcohol dependence disorders (32) and Parkinson’s disease (33). Chromosome 11 has been linked to the development of ADHD, autism, depression, schizophrenia and mental disorders in general (34, 35). Chromosome 3 has also been linked to depression, drug dependence (36), schizophrenia and bipolar disorder (37). Lastly, chromosome 2 has been linked to the development of autism, Dementia and Lewy’s body (38, 39), Parkinson’s and Alzheimer’s disease (40, 41)

Finally, it is worth noting that the majority of the most “culpable” genetic targets for the occurrence of these disorders in our study belong to the superfamily of GPCRs (DRD1, DRD2, DRD3, DRD4, HTR1B, OXTR, OPRM1 etc.) and to neurotrophic factors (BDNF, BDNF-AS). Research interest recently has been aimed at GPCR proteins, as they themselves seem to be risk factors but also future therapeutic targets for many disorders and diseases, including mental health disorders (42). Especially for the Dopamine Receptors D, there are numerous studies which link them to behavioral disorders, through DRD4 and its 7-repeat allele polymorphism (13). DRD4 polymorphisms in general have been studied extensively in relation to behavioral disorders and are now potential biomarkers for individualized therapeutic approaches (14). As for the neurotrophic factors, they also have been significantly associated with the occurrence of cognitive and behavioral disorders, with the main representative of the risk of the latter, BDNF. Neurotrophins play an important role in the development, differentiation, maintenance and survival of distinct and overlapping neuronal populations in the central and peripheral nervous systems(16). Recent studies have linked BDNF to the pathophysiology of cognitive, psychiatric, and neurodegenerative diseases with a possible mechanism possibly due to a nucleotide polymorphism induced by the change of valine (Val) to methionine (Met) at position 66 in the proteome BDNF. Among other diseases, depression, bipolar disorder, schizophrenia are the main mental / behavioral disorders associated with this polymorphism of the *BDNF* gene (16). Particularly, in our study, only BDNF showed evidence of significant association with cognition, in addition to DRDs that showed also significant association with behavior. The most interesting point of these results is that BDNF was also found here to be associated with behavior as well. Also, various studies have supported our findings about BDNF in cognition and behavior(16, 43-45).

Our analysis indicates a potential linkage of DRDs and BDNF through the dopaminergic system that can lead to an emergence of the disorders studied. Dopamine has been found to be a hormone that affects the development of these disorders by fluctuating its levels (13). A possible decrease in BDNF secretion may also lead to decreased dopamine levels, which signal the onset of mental disorders (43). DRDs additionally regulate the basic expression of dopamine levels and these receptors can also activate the BDNF factor, whose expression is controlled by its receptor, TrkB (46). We understand, therefore, that if a malfunction or mutation or polymorphism occurs in any of the above candidate genetic factors, dopamine can be affected and potentially contribute to the emergence of the disorders. More specifically, malfunction of the dopamine receptors can lead to development of a behavioral disorder, while malfunction in BDNF can lead to development of a cognitive disorder (12-14). Comorbidity can also occur if the system is at its most dysregulated.

## Conclusions

Clinical genomics has emerged to be a great scientific implement, which Geneticists and Bioinformaticians can use, through its tools and applications, in order to prevent, diagnose and treat diseases and disorders, in concert with medical doctors and other scientists, under the aegis of Personalized Medicine. In the present study and through a novel semantic analysis pipeline, we managed to correlate and highlight the significance of the dopaminergic system. More precisely, we linked Dopamine Receptors D (DRDs) to Brain Derived Neurotrophic Factor in the occurrence of cognitive and behavioral disorders. However, these disorders are due to a combination of genetic prevalence and epigenetic factors. Several studies have pointed out that the most positive results, in terms of a holistic confrontation of this medical issue, come from a combination of pharmacological treatment and psychotherapeutic techniques, like Cognitive-Behavioral Therapy (CBT) (47). Because stress is, among other things, perhaps the most important epigenetic factor for the occurrence of these disorders, CBT can help patients reduce the impact of such factors in their lives, and therefore reduce the effect of the symptoms of the disorders. Along with medication and CBT, researchers have highlighted the positive role of physical exercise on the lives of people suffering from such a disorder, as well as the proper healthy nutrition and sufficient sleep (48), (49). Based on the results of our genetic study, many studies have also shown that exercise and a healthy diet program can maintain dopamine to normal levels, thus preventing their dysfunction, by regulating the levels of BDNF and DRDs through the dopaminergic system. Nevertheless, validation of these findings is required before firm conclusions can be made, and clinical or pharmacological utility can be appraised.

## Acknowledgments

Not applicable.

